# Arabidopsis GLK transcription factors interact with ABI4 to modulate cotyledon greening in light-exposed etiolated seedlings

**DOI:** 10.64898/2026.01.22.701071

**Authors:** Pengxin Yu, Friederike Saga, Miriam Bäumers, Ute Hoecker

## Abstract

During seedling etiolation in darkness, the biosynthesis of protochlorophyllide (Pchlide) and the development of etioplasts must be strictly controlled to prevent photooxidative damage upon light exposure. The transcription factors GLK1 and GLK2 are central regulators of chlorophyll biosynthesis and chloroplast biogenesis. Here, we show that GLK1 and GLK2 interact with ABSCISIC ACID INSENSITIVE 4 (ABI4). We reveal that GLKs and ABI4 have antagonistic functions in cotyledon greening of light-exposed etiolated seedlings: compared to the wild type, *abi4* mutants, similar to a transgenic line overexpressing GLK2, accumulated more Pchlide in dark-grown seedlings, while *glk1 glk2* mutants contained less Pchlide. The high Pchlide levels in etiolated *abi4* mutants and GLK2 overexpressors were inefficiently photoreduced upon light exposure, leading to a significant accumulation of ^1^O_2_ in the cotyledons after the dark-to-light transition. This corresponded to low cotyledon greening rates and low seedling survival. Additionally, we identified eight *PhANG*s involved in Pchlide biosynthesis and etioplast development, whose transcript accumulation patterns may contribute to the photobleaching of etiolated *abi4* mutants and GLK2 overexpressors. Importantly, the high Pchlide content, low cotyledon greening rate, high ^1^O_2_ level and high *PhANG* induction in *abi4* mutant seedlings were fully dependent on *GLK1* and *GLK2*, indicating that *ABI4* acts upstream of *GLKs.* Since ABI4 does not regulate *GLK* transcript level, and ABI4 physically interacts with GLK proteins, these data suggest that ABI4 inhibits GLK1 and GLK2 activities in etiolated seedlings to prevent high Pchlide accumulation which would lead to high ^1^O_2_ levels and seedling death upon exposure to light.

**Significance statement:** The transition from darkness to light is a critical moment that determines seedling establishment and survival. We report here that the transcription factor ABI4 inhibits GLK1 and GLK2 activities during seedling etiolation to repress Pchlide biosynthesis in darkness, thereby limiting singlet oxygen accumulation and seedling death upon exposure to light.

## Introduction

After germination, plant seedlings are faced with the need to adjust their developmental programs in response to specific light conditions. If seedlings germinate under the soil, they will grow in darkness in a process termed etiolation (or skotomorphogenesis), during which specialised chloroplast precursors, known as etioplasts, initiate their development. Within the etioplasts, the enzyme protochlorophyllide reductase (POR) accumulates along with the chlorophyll precursor protochlorophyllide (Pchlide). In angiosperms, PORs are responsible for the only light-dependent step in chlorophyll biosynthesis, and this step occurs in a para-crystalline etioplast compartment called prolamellar body (PLB), where Pchlide forms a ternary complex with PORs and nicotinamide adenine dinucleotide phosphate (NADPH). Pchlide in Arabidopsis can be divided into two categories: the photoactive Pchlide is bound by PORs, whereas the non-photoactive Pchlide remains unbound during etiolation (Klein and Schiff, 1972; Lebedev and Timko, 1999; Li *et al*., 2025).

Etiolated seedlings grow upwards through the soil by sensing gravity (Kawamoto and Morita, 2022; Kim *et al*., 2011), and when they reach the soil surface they are immediately exposed to light. This is a critical moment that determines plant survival and health, during which the seedlings respond to light in a process termed de-etiolation or photomorphogenesis. Two key photomorphogenic traits are chlorophyll biosynthesis and chloroplast biogenesis (overall termed greening), both being crucial for the establishment of photoautotrophic growth. Chlorophyll biosynthesis refers to the enzymatic processes where the PORs convert photoactive Pchlide into chlorophyllide (Chlide) in a reaction termed photoreduction, and Chlide is subsequently converted into chlorophyll (Apel *et al*., 1980; Armstrong *et al*., 1995). In contrast, chloroplast biogenesis is represented by structural changes in the plastid, where etioplasts differentiate into chloroplasts by expanding the thylakoid surface area while integrating and arranging photosynthetic complexes into the thylakoids (Pipitone *et al*., 2021). Chlorophyll is synthesized in Arabidopsis cotyledons within the first four hours of illumination, while the assembly of a fully photosynthetic chloroplast takes more than ten hours during de-etiolation (Pipitone *et al*., 2021).

The process of greening requires a balancing act. Both chlorophyll biosynthesis and chloroplast biogenesis are needed for the maximization of photosynthetic rate. Consequently, having a low chlorophyll content and less differentiated chloroplasts can both reduce photosynthetic performance and yield (Frangedakis *et al*., 2024; Kim *et al*., 2023). On the other hand, non-photoactive Pchlide can absorb light energy (termed photosensitization) and transfer this energy to atmospheric oxygen, generating potentially toxic molecules called reactive oxygen species (ROS). As a result, excessive Pchlide accumulation during etiolation can lead to seedling lethality by producing ROS during dark-to-light transitions (Meskauskiene *et al*., 2001). Therefore, plastid activity needs to be appropriately repressed in darkness and activated in light for an optimized plant development.

Plastid development and activity depend on the transcriptional states of photosynthesis-associated nuclear genes (*PhANG*s), because over 90% of chloroplastic proteins are nucleus-encoded (Martin *et al*., 2002). Multiple classes of transcription factors form a network to tightly regulate the transcription of *PhANGs*. In the dark, PHYTOCHROME-INTERACTING FACTORs (PIFs) act as a major class of *PhANG* repressors (Huq *et al*., 2004; Stephenson *et al*., 2009; Monte *et al*., 2004). PIFs repress etioplast development and Pchlide biosynthesis in concert with ETHYLENE-INSENSITIVE3 (EIN3) and ETHYLENE-INSENSITIVE3-LIKE1 (EIL1) (Liu *et al*., 2018; Zhong *et al*., 2009). In the light, ELONGATED HYPOCOTYL5 (HY5), GATA NITRATE-INDUCIBLE CARBON-METABOLISM-INVOLVED (GNC), CYTOKININ-RESPONSIVE GATA1 (CGA1), MYB-related transcription factors (MYBS) and GOLDEN2-LIKE (GLK) promote greening by activating *PhANG* transcription either alone or cooperatively (Oyama *et al*., 1997; Chiang *et al*., 2012; Frangedakis *et al*., 2024; Fitter *et al*., 2002; Zhang *et al*., 2024).

GOLDEN2-LIKE 1 and 2 (GLK1 & 2) are key greening regulators that have led to significant agricultural applications, where ectopic expression of maize GLKs improved yield in rice by 30-70% (Li *et al*., 2020; Yeh *et al*., 2022). GLKs exist in most plants as a paralogous pair (Fitter *et al*., 2002; Powell *et al*., 2012; Tu *et al*., 2022; Rossini *et al*., 2001), and they mainly act in the nuclei of mesophyll cells (Waters *et al*., 2008), where they promote the transcription of the *PhANG*s encoding thylakoid-associated protein components and chlorophyll biosynthetic enzymes (Waters *et al*., 2009; Tu *et al*., 2022). The necessity of fine-tuning GLK activity was demonstrated by M. Li et al. (2022), in which two transcription factors were reported to physically and antagonistically interact with GLKs, ultimately modulating cell death in response to external stress. Although *GLK* transcript levels are low in darkness (Fitter *et al*., 2002), and GLK protein is degraded in darkness (D., Zhang *et al*., 2021), GLKs still play a physiological role in seedling etiolation. Specifically, Arabidopsis *glk1 glk2* mutant seedlings accumulate less Pchlide and contain smaller etioplasts with smaller PLBs when compared to the wild type (Waters *et al*., 2009). Therefore, maintaining a moderate level of GLK activity during etiolation may also be vital for seedling health. Since multiple studies have demonstrated the physiological significance of the post-translational regulation of GLKs in different plant species (Han *et al*., 2024; Y., Li *et al*., 2022; Ni *et al*., 2017; Rauf *et al*., 2013; Tachibana *et al*., 2024; Tang *et al*., 2016; C., Zhang *et al*., 2021; D., Zhang *et al*., 2021; Zhang *et al*., 2024; Sun *et al*., 2025), we screened for other transcription factors that physically interact with GLK1 or GLK2 from Arabidopsis.

## Results

### ABI4 and GLKs interact in yeast two-hybrid and *in planta*

To identify proteins interacting with GLKs, we screened an Arabidopsis transcription factor library (Paz-Ares *et al*., 2002) using a yeast two-hybrid workflow devised by Castrillo et al. (2011). In order to use GLKs as bait, we first deleted the transactivation domain from the GLKs, then performed the screen using the GLK C-terminal domains as baits (Figure 1a). We identified an interaction between ABSCISIC ACID INSENSITIVE 4 (ABI4) and the C-terminal domain of GLK2 (Figure 1b).

**Figure 1.**
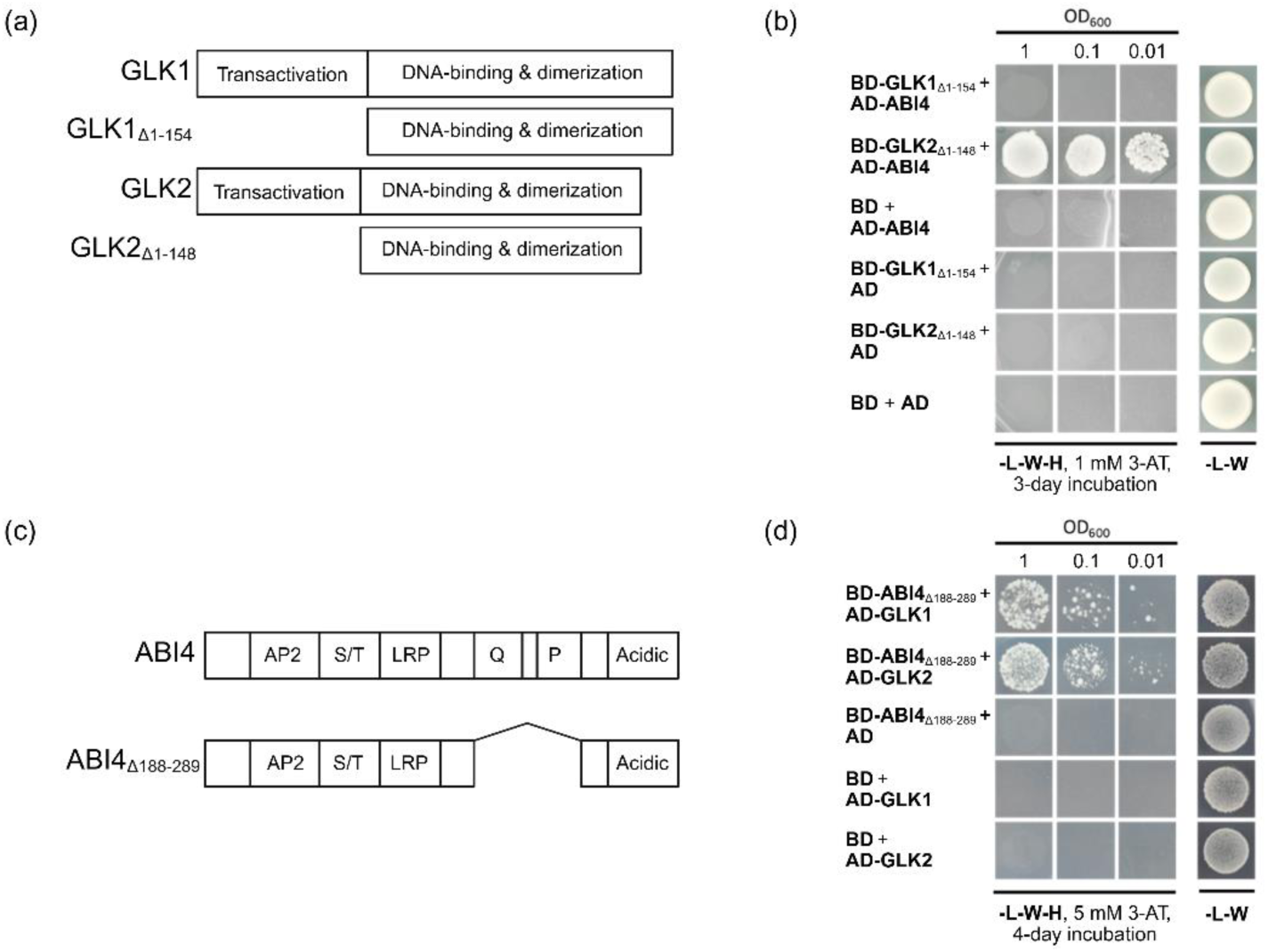
GLK1 and GLK2 physically interact with ABI4 in yeast two-hybrid, **(a)** Scheme of GLK1 and GLK2 proteins and deletion-derivatives lacking the transactivation domain. The latter were used as baits in the yeast two-hybrid screen, **(b)** GLK2_Δ1-154_ interacts with ABI4. **(c)** Scheme of ABI4 protein and a deletion-derivative lacking the transactivation domain, the latter was used as the bait in the following yeast two-hybrid assay. AP2 = APETALA2 domain; S/T = serine-/threonine-rich domain; LRP = LRP motif; Q and P = glutamine- and proline-rich domains, respectively, **(d)** ABI4_Δ188-28g_ interacts with full-length GLK1 and GLK2.

Due to the sharing of conserved regions and functional redundancy between GLK1 and GLK2 (Rauf et al., 2013; Waters et al., 2009), we hypothesized that ABI4 interacts with GLK1 as well as GLK2. Although the GLK1 C-terminal domain did not interact with ABI4 (Figure 1b), we hypothesized that GLK1 N-terminal domains may be necessary for an interaction between GLK1 and ABI4. To test this, we deleted the transactivating glutamine-rich and proline-rich domains from ABI4 so that we can use ABI4 as a bait (Figure 1c). Indeed, ABI4_Δ188-289_ interacted with full-length GLK1 and GLK2 in yeast two-hybrid (Figure 1d).

We further confirmed the ABI4–GLK interaction *in planta*. First, we performed Förster resonance energy transfer by fluorescence lifetime imaging (FRET-FLIM) experiments after co-expressing YFP-tagged ABI4 and mCherry-tagged GLK proteins in leek epidermal cells by particle bombardment. As shown in Figure 2a and 2b, the lifetime of the donor fluorophore YFP was significantly reduced when YFP-ABI4 was co-expressed with either mCherry-GLK1 or mCherry-GLK2 compared to co-expression with mCherry, supporting an interaction between Arabidopsis ABI4 and GLK1 as well as GLK2. Next, we performed a luciferase complementation imaging assay after co-expressing nLUC- and cLUC-fused proteins in tobacco by *Agrobacterium* infiltration. We observed that both GLK1-nLUC and GLK2-nLUC generated luminescence when co-expressed with cLUC-ABI4 (Figure 2c and 2d), whereas the negative controls did not generate luminescence, supporting an interaction between Arabidopsis ABI4 and GLK in tobacco. Together, these assays corroborate the existence of an *in-planta* interaction between ABI4 and both GLKs.

**Figure 2.**
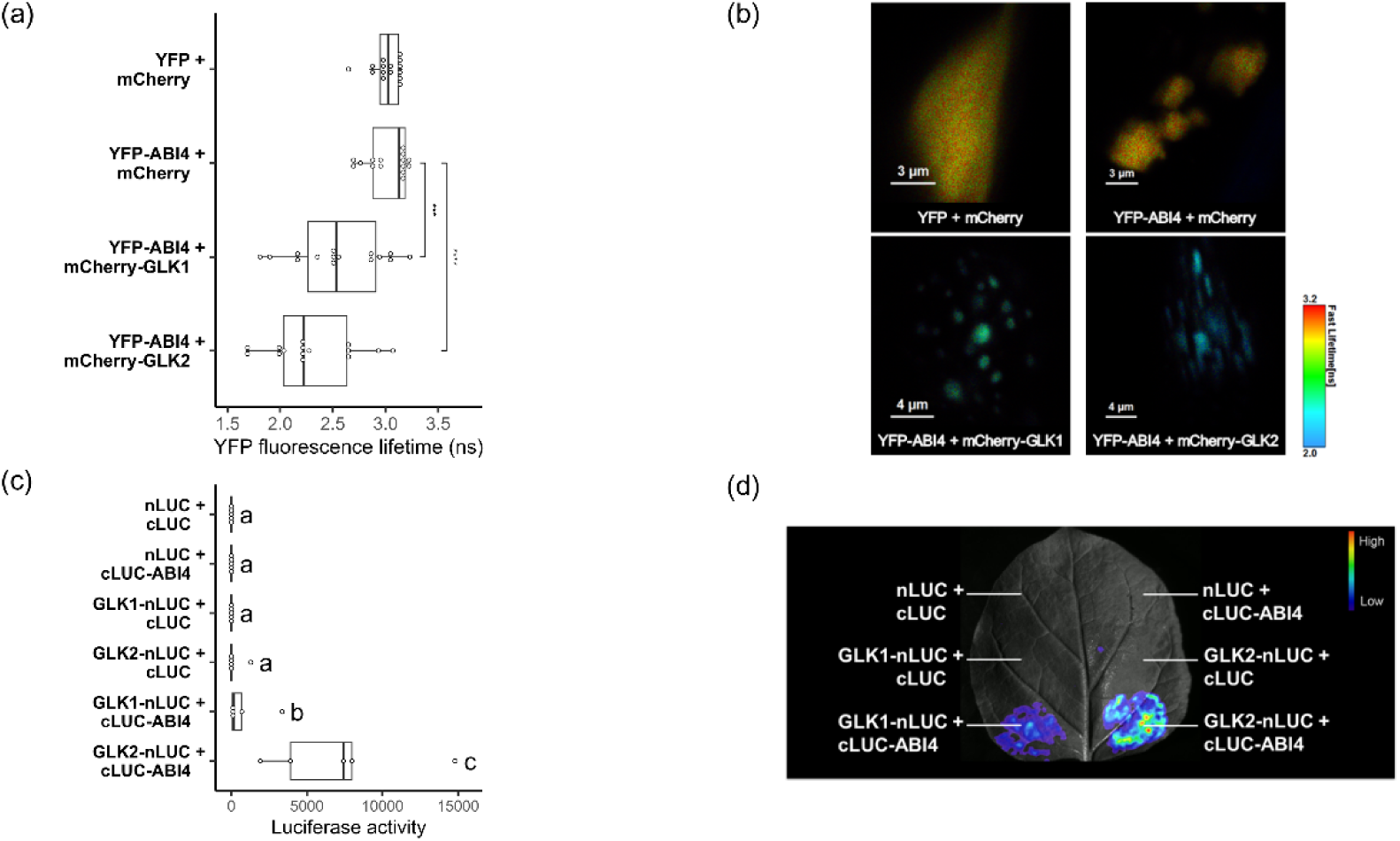
GLK1 and GLK2 physically interact with ABI4 *in planta,* **(a)** Lifetime of donor (YFP) fluorophore measured by FRET-FLIM inside the nuclei of leek epidermal cells after particle bombardment, n=15 nuclei. Asterisks indicate a significant difference in lifetime between the indicated pair (Student’s t-test, ***P< 0.001, ****P < 0.0001). **(b)** YFP FLIM images of the representative nuclei from (a) showing changes in lifetime, **(c)** Quantification of luminescence detected by a CCD camera from a luciferase complementation imaging assay in tobacco leaves, n=5 leaves. Combinations which are significantly different (P < 0.05) are labelled with different letters (Pairwise Wilcoxon test with multiple testing correction), **(d)** Representative leaf image from (c) showing the luminescence signal in sectors transfected with the indicated plasmid combinations.

### Etiolated *abi4* mutant seedlings display *GLK*-dependent Pchlide overaccumulation and inefficient Pchlide photoreduction

The ABI4-GLK interaction prompted us to investigate the biological relevance of this interaction. To this end, we compared the phenotypes of *abi4* and *glk1 glk2* loss-of-function mutants. First, because GLKs promote chlorophyll accumulation in light (Fitter *et al*., 2002), we quantified the chlorophyll content in seedlings grown on soil in a long-day photoperiod for 10 days. As described previously (Waters *et al*., 2008), the *glk1 glk2* double mutant accumulated significantly less chlorophyll than the *Col-0* wild type, while the *35Spro*::*GLK2*-*GFP* line accumulated slightly more chlorophyll than the wild type, though the difference was not statistically significant (Figure S1). We included two *abi4* knock-out mutant alleles: *abi4-1*, with a 1-bp deletion at *ABI4* codon 157 in the *Col-0* background (Finkelstein *et al*., 1998) and *abi4-103 gl1*, with a nonsense point mutation at *ABI4* codon 39 in the *gl1* background (Laby *et al*., 2000). Both *abi4* mutants accumulated similar levels of chlorophyll compared with their wild-type genotypes, indicating that the ABI4-GLK interaction does not play a role in chlorophyll homeostasis under these conditions.

Next, we hypothesized that the ABI4-GLK interaction plays a specific role in Pchlide accumulation during seedling skotomorphogenesis. In the following phenotypic assays, we included two *abi4* knock-out mutant alleles: the above-mentioned *abi4-1*, and *abi4-t* which contains a T-DNA insertion in *ABI4* (Shu *et al*., 2013; Alonso *et al*., 2003). To elucidate the genetic relationships between *ABI4* and *GLK*s, we generated the *glk1 glk2 abi4-1* triple mutant by crossing *glk1 glk2* with *abi4-1*. The *abi4-1* homozygosity in the *glk1 glk2 abi4-1* triple mutant was verified by genotyping using a derived Cleaved Amplified Polymorphic Sequences (dCAPS) marker (Figure S2a) and observing a complete insensitivity to 1.5 µM ABA during seed germination (Figure S2b). The *glk1 glk2* homozygosity in the *glk1 glk2 abi4-1* triple mutant was confirmed by the observation that the triple mutant has the same chlorophyll-deficient phenotype as the *glk1 glk2* double mutant (Figure 4a).

**Figure 3.**
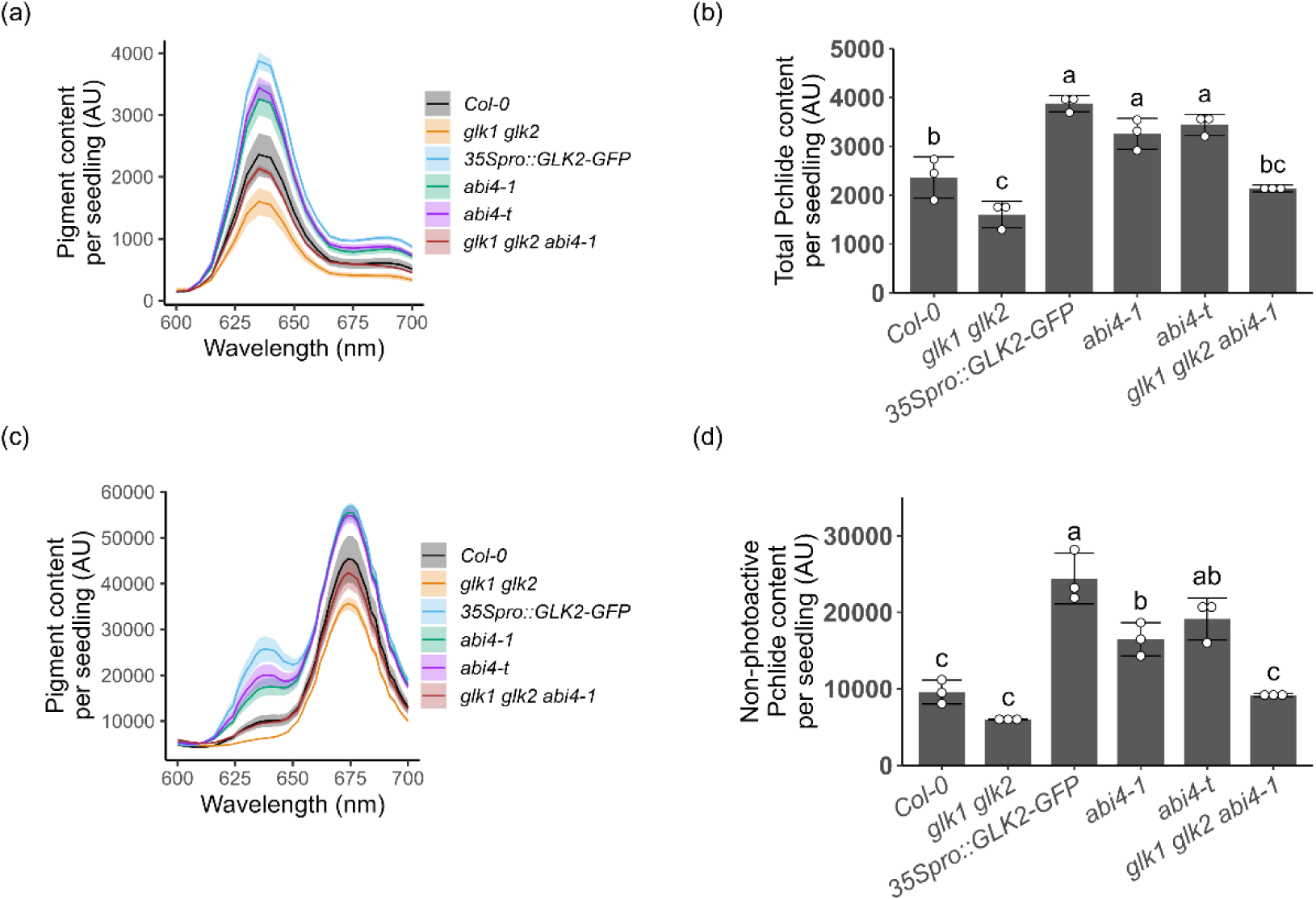
Etiolated *abi4* mutants overaccumulate Pchlide and cannot efficiently convert all Pchlide into Chlide, and these phenotypes are dependent on *GLK1* and *GLK2.* **(a)** Fluorescence of Pchlide at different emission wavelengths in 5-day-old etiolated seedlings, data are plotted as mean (solid lines) ± SD (shaded areas), (b) Comparison of total Pchlide content by the peak fluorescence at 634 nm from (a). Genotypes which are significantly different are labelled with different letters (n=3 replicate extracts, ANOVA and Tukey’s post hoc tests), error bars indicate SD. **(c)** Fluorescence of Pchlide (634 nm) and Chlide (670 nm) in 5-day-old etiolated seedlings after transfer from darkness to 100 µmol m^−2^ s^−1^ white light for 5 min. **(d)** Comparison of non-photoactive Pchlide content using the peak fluorescence at 634 nm from (c). Genotypes which are significantly different are labelled with different letters (n=3 replicate extracts, ANOVA and Tukey’s post hoc tests), error bars indicate SD.

**Figure 4.**
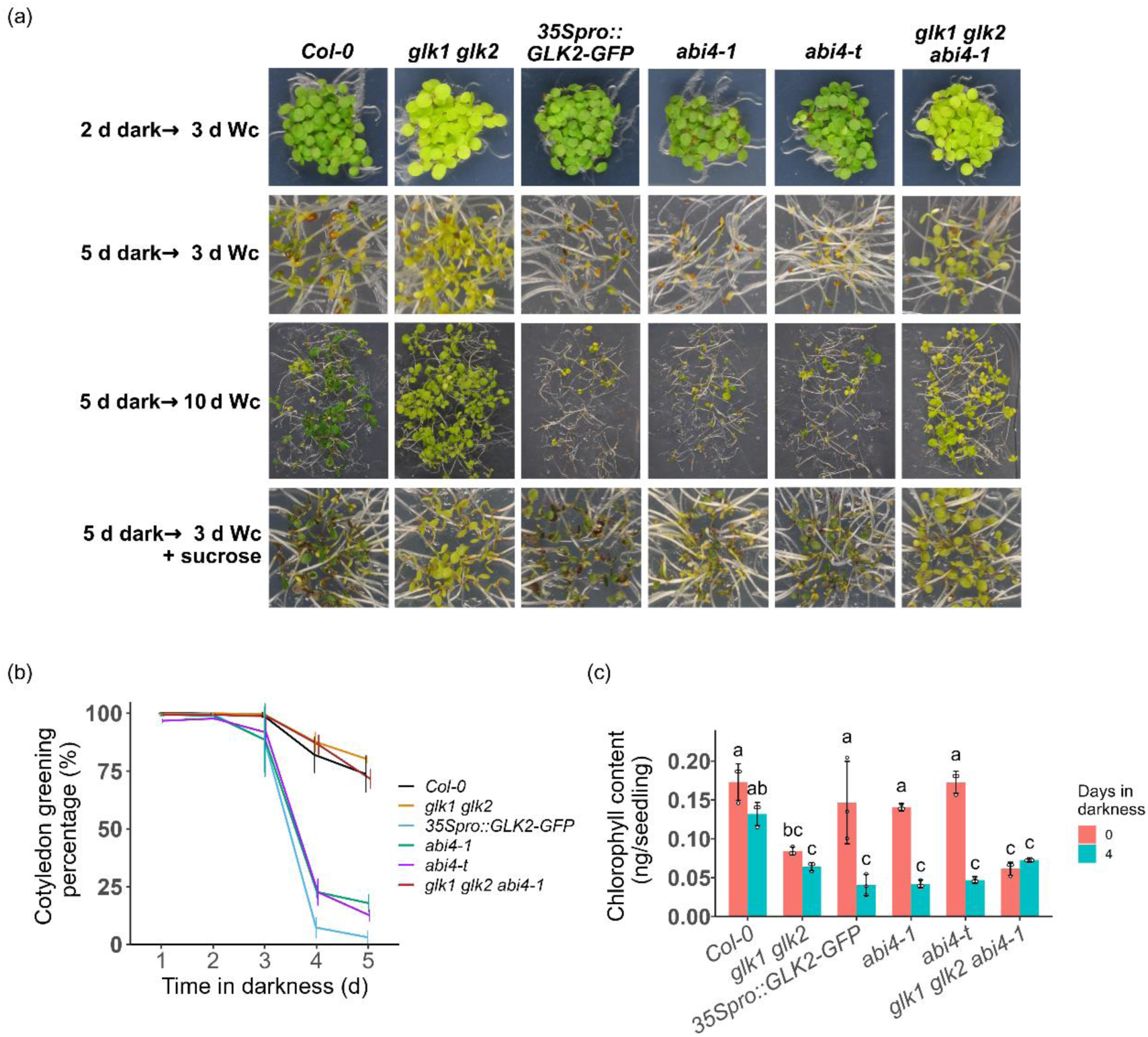
A *35Spro::GLK2-GFP* line and two *abi4* mutants fail to green when seedlings etiolated for more than 4 days are exposed to light, **(a)** Photographs of seedlings that were etiolated for the indicated number of days followed by transfer to constant white light (Wc) for the indicated time. Seedlings shown in the last row were grown on 2% sucrose, **(b)** Quantification of cotyledon greening percentage of etiolated seedlings exposed to Wc for 3 days, n=3 replicate plates, error bars indicate SD. **(c)** Quantification of chlorophyll content of seedlings grown either without prior etiolation or with 4 days of etiolation, then exposed to Wc for 5 days. Genotypes which are significantly different are labelled with different letters (n=3 replicate extracts, ANOVA and Tukey’s post hoc tests), error bars indicate SD.

To determine Pchlide levels, we extracted and quantified the total Pchlide content in 5-day-old etiolated seedlings. We found that in comparison to the wild type, the *glk1 glk2* mutant accumulated significantly less Pchlide, while the *35Spro*::*GLK2*-*GFP* line accumulated more Pchlide (Figure 3a and 3b). Importantly, both *abi4* mutants had higher levels of Pchlide, thus resembling the *35Spro*::*GLK2*-*GFP* line, while the Pchlide content returned to a wild-type level in the *glk1 glk2 abi4-1* mutant (Figure 3a and 3b). This revealed that the total Pchlide overaccumulation in etiolated *abi4* mutants was completely dependent on *GLK*s.

According to previous studies, photoactive Pchlide is converted by PORs to Chlide by one millisecond of white light, while non-photoactive Pchlide is not efficiently photoreduced (Shibata, 1957; Xu *et al*., 2016). Therefore, we measured the non-photoactive Pchlide levels in 5-day-old etiolated seedlings exposed to 5 min of white light. We found that, similar to the *35Spro*::*GLK2*-*GFP* line, both *abi4* mutants did not efficiently convert all Pchlide to Chlide (Figure 3c), because they still contained more non-photoactive Pchlide than the wild type upon light exposure (Figure 3d). In contrast, all Pchlide was photoreduced in both the *glk1 glk2* mutant and the *glk1 glk2 abi4-1* mutant (Figure 3c), where the non-photoactive Pchlide content returned to a wild-type level (Figure 3d). We conclude that etiolated *abi4* mutant seedlings display inefficient Pchlide photoreduction and non-photoactive Pchlide overaccumulation, and that these phenotypes are completely dependent on *GLK*s.

### Etiolated *abi4* mutant seedlings exhibit a *GLK*-dependent reduced greening rate when exposed to light

Non-photoactive Pchlide is a photosensitizer that can cause pigment photobleaching and inhibition of greening (Erdei *et al*., 2005; Meskauskiene *et al*., 2001). Since etiolated *abi4* mutant seedlings contained more non-photoactive Pchlide compared to the wild type, we performed cotyledon greening assays using seedlings that were first etiolated for 1 to 5 days and then exposed to 3 days of white light. A visual examination revealed that 2-day-old etiolated seedlings of all investigated genotypes developed green cotyledons after transfer to white light (Figure 4a, first row). However, when seedlings were etiolated for 5 days before the transfer to white light (Figure 4a, second row), a small portion of the *Col*-*0* wild-type seedlings failed to develop green cotyledons. In comparison, most seedlings of the *35Spro*::*GLK2*-*GFP* line and the two *abi4* mutants had cotyledons that failed to green, and this low greening rate was rescued in the *glk1 glk2 abi4*-*1* triple mutant. After prolonging the light exposure to 10 days (Figure 4a, third row), we observed that higher cotyledon greening rates corresponded to higher seedling establishment and survival rates, while seedlings that failed to green showed arrested development. We therefore designate the fail-to-green cotyledon phenotype as a photobleaching phenotype. Since sucrose is known to suppress cotyledon photobleaching by stimulating Pchlide photoreduction (Barnes *et al*., 1996; Mu *et al*., 2022), we assessed the effect of adding 2% sucrose exogenously to the culture medium. As shown in Figure 4a (fourth row), sucrose indeed partially rescued cotyledon photobleaching, especially in the *35Spro*::*GLK2*-*GFP* line and the two *abi4* mutants, suggesting that the photobleaching phenotype in these lines could be due to inefficient Pchlide photoreduction.

We quantified the proportion of greened seedlings by visually categorizing seedlings as green or bleached. As seen in Figure 4b, the *35Spro*::*GLK2*-*GFP* line and the two *abi4* mutants displayed consistently lower greening rates than *Col*-*0* and *glk1 glk2* when etiolation lasted longer than 3 days. Importantly, the *glk1 glk2* mutations in the *abi4*-*1* background rescued the failure of *abi4-1* to green. To further quantify the degree of photobleaching, we measured the chlorophyll levels in seedlings grown either without etiolation or with 4 days of etiolation prior to exposure to white light for 5 days. As shown in Figure 4c, in seedlings grown without prior etiolation, *Col*-*0*, *35Spro*::*GLK2*-*GFP* and the two *abi4* mutants contained similar levels of chlorophyll, whereas the chlorophyll levels of *glk1 glk2* and *glk1 glk2 abi4*-*1* mutants were similar to each other, but significantly lower than that of the wild type. The 4 days of prior etiolation did not significantly change the light-induced chlorophyll accumulation level in *Col*-*0* nor *glk1 glk2* mutant seedlings, but it significantly decreased the seedling chlorophyll levels in *35Spro*::*GLK2*-*GFP* and the two *abi4* mutants. Strikingly, the *glk1 glk2* mutations rescued the inhibited chlorophyll accumulation caused by the 4 days of prior etiolation in the *abi4-1* mutant background, demonstrating again that the *GLK*s are necessary for the photobleaching of light-exposed etiolated seedlings. To summarize, etiolated *abi4* mutants and *35Spro*::*GLK2*-*GFP* seedlings have lower greening rates compared to the wild type when exposed to light, and this photobleaching phenotype is dependent on *GLK*s.

### Photobleaching of the etiolated seedlings corresponds to singlet oxygen accumulation in the cotyledons

Although multiple species of ROS exist in Arabidopsis, the cause of Pchlide-induced photobleaching has been attributed to the generation of singlet oxygen (^1^O_2_) during photosensitization (Meskauskiene et al., 2001; Triantaphylidès et al., 2008). Hence, we quantified the ^1^O_2_ generated in 5-day-old etiolated seedlings before and after exposure to light. To detect ^1^O_2_ *in planta*, we stained seedlings with the fluorescent probe Singlet Oxygen Sensor Green (SOSG) (Flors *et al*., 2006) and then detected both SOSG fluorescence and chlorophyll autofluorescence in the cotyledons using a confocal microscope. Before exposure to white light (0 d Wc), all genotypes contained a low level of ^1^O_2_ and displayed limited chlorophyll autofluorescence (Figure 5a, b). After 3 days of light exposure (3 d Wc), the *35Spro::GLK2-GFP* line and both *abi4* mutants failed to accumulate chlorophyll, and this phenotype correlated with high ^1^O_2_ accumulation levels. In contrast, light-exposed wild-type seedlings and—to a lesser extent—*glk1 glk2* and *glk1 glk2 abi4*-*1* mutant seedlings exhibited chlorophyll autofluorescence and low levels of ^1^O_2_. Together, these ^1^O_2_ levels are consistent with the Pchlide and photobleaching phenotypes reported above: when compared to the wild type, etiolated *abi4* mutant and *35Spro::GLK2-GFP* seedlings contained higher levels of non-photoactive Pchlide, displayed low greening rates and high ^1^O_2_ content when exposed to light. Again, the lower ^1^O_2_ level generated by photosensitization in the *glk1 glk2 abi4-1* mutant seedlings compared to that in the *abi4-1* mutant seedlings supports the conclusion that the photobleaching observed in etiolated *abi4* mutant seedlings is dependent on GLKs, and that GLK activity plays a central role in singlet oxygen accumulation when etiolated seedings are exposed to light.

**Figure 5.**
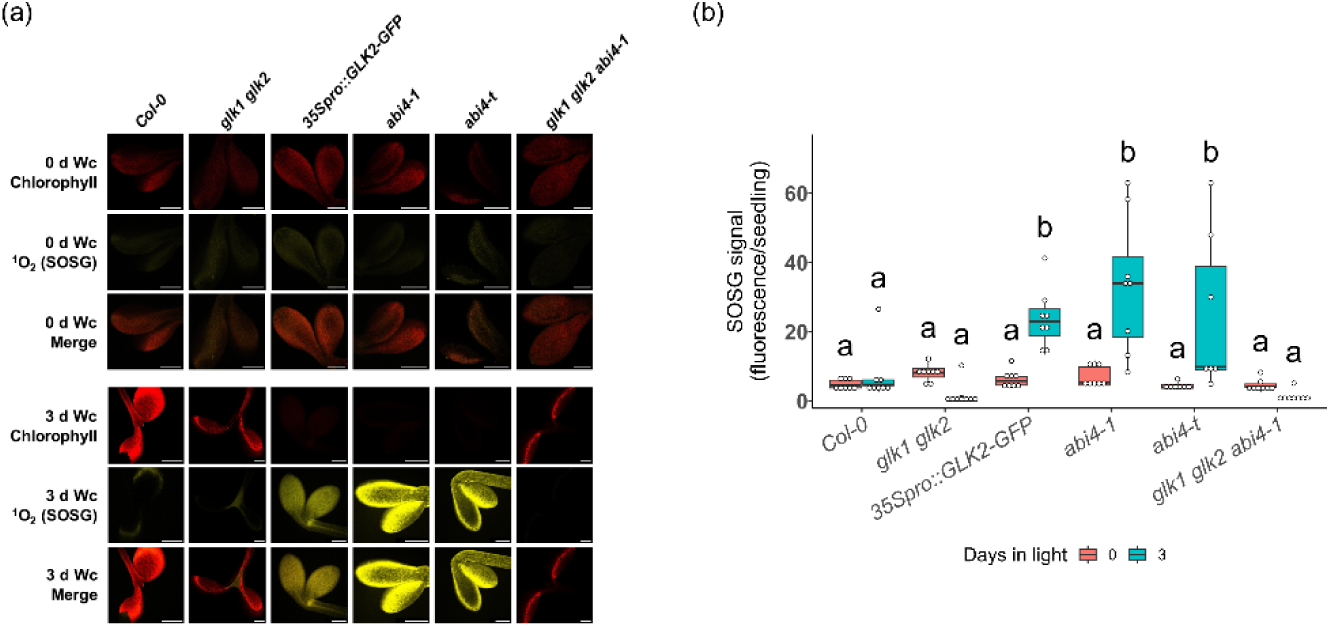
Photobleaching of etiolated, then light-exposed *abi4* mutant and *35Spro:.GLK2- GFP*seedlings correlates with high singlet oxygen (^1^O_2_) accumulation, **(a)** Representative fluorescence microscopy images of chlorophyll autofluorescence and Singlet Oxygen Sensor Green fluorescence (SOSG, detecting ^1^O_2_) in 5-day-old etiolated seedlings before (0 d Wc) and after exposure to 100 µmol m^-2^ s^-1^ constant white light for 3 days (3 d Wc). Scale bars = 200 µm. **(b)** Quantification of SOSG fluorescence per seedling in cotyledons of seedlings grown as in (a). Groups which are significantly different are labelled with different letters (n=8 seedlings, ANOVA and Tukey’s post hoc tests).

### Certain *PhANG* transcripts show GLK-dependent upregulation in etiolated *abi4* mutant seedlings

We established above that the photobleaching of etiolated *abi4* mutant seedlings is likely due to an altered Pchlide homeostasis. Because *PhANG*s encode etioplast proteins responsible for Pchlide biosynthesis and photoreduction (Solymosi and Aronsson, 2013), and GLKs are transcriptional activators of *PhANG*s (Waters *et al*., 2009), we first hypothesized that *GLK* transcript levels may be altered in the *abi4* mutants. To test this, we quantified the transcript levels of *GLK1* and *GLK2* by performing RT-qPCR using RNA isolated from 5-day-old etiolated seedlings. However, as shown in Figure 6a, both *abi4* mutants showed similar *GLK1* and *GLK2* transcript levels to those in the wild type, indicating that *ABI4* did not affect *GLK1* nor *GLK2* transcript levels under these conditions. There were also no significant changes observed for *GLK1* nor *GLK2* transcript levels in the *glk1 glk2 abi4-1* mutant compared to the *glk1 glk2* mutant, indicating that *ABI4* did not affect *GLK1* nor *GLK2* transcript levels in this genetic background either. Thus, the enhanced photobleaching observed in *abi4* mutants is not caused by a change in *GLK1* or *GLK2* transcript levels.

**Figure 6.**
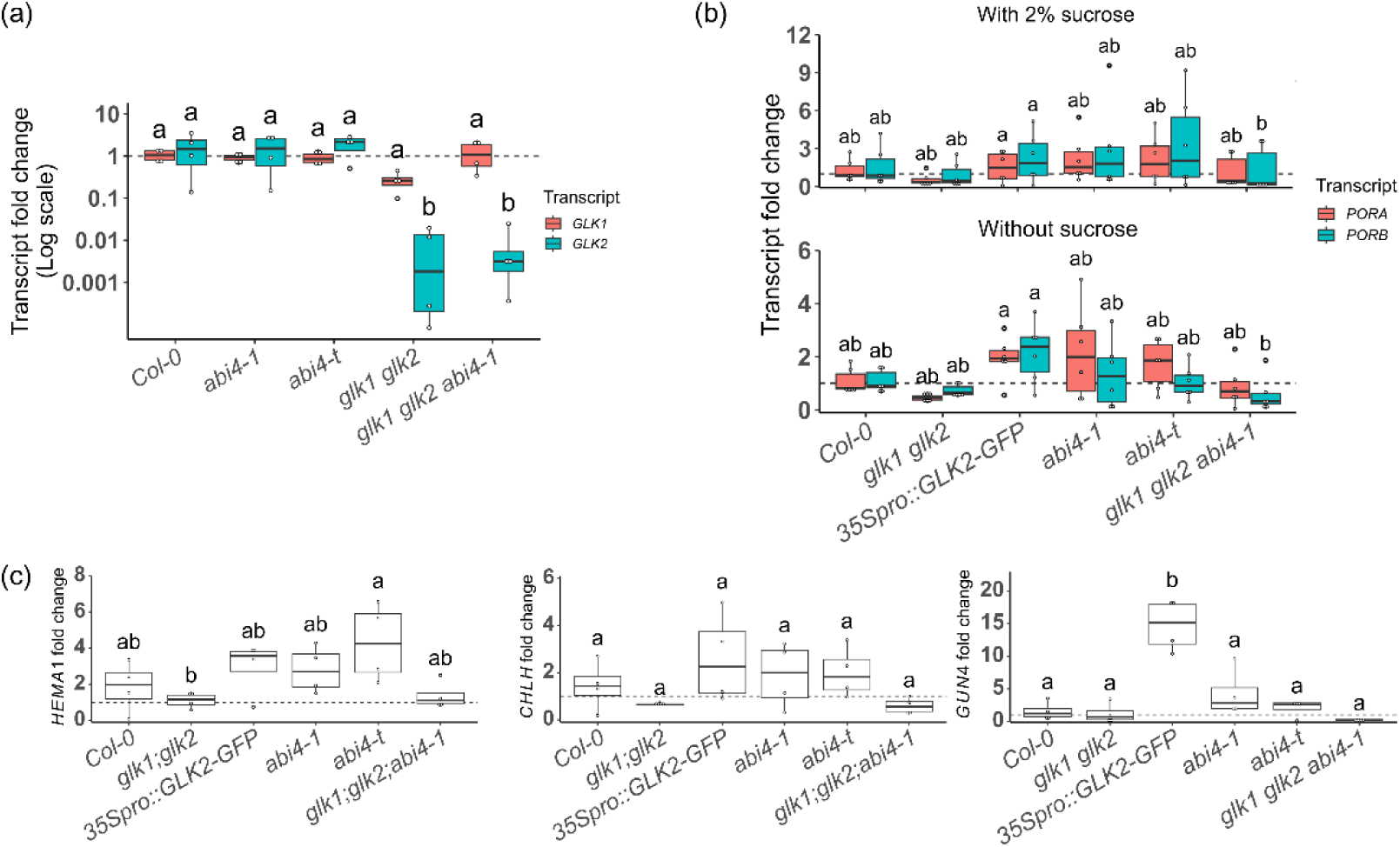
Transcript fold change of *GLKs* and Pchlide biosynthesis genes in 5-day-old etiolated seedlings. Seedlings were grown on MS agar plates without sucrose (unless indicated otherwise), and the transcript levels were quantified using RT-qPCR. Mean transcript level in *Col-0* was set to 1 (indicated by the dashed line), and data in each panel were compared with ANOVA and Tukey’s post hoc tests. Genotypes which are significantly different are labelled with different letters, **(a)** Transcript levels of *GLK1* and *GLK2* are not altered in *abi4* mutants, n=4. **(b)** Transcript levels of *PORA* and *PORB* are not lower in *abi4* mutants with (top panel) or without 2% sucrose (bottom panel), n-6. **(c)** Three main Pchlide biosynthesis genes are targeted by GLKs (n=4 replicate extracts).

Next, we hypothesized that low *POR* transcript accumulation also contributed to the inefficient Pchlide photoreduction in *abi4* mutant seedlings. Since sucrose rescued the photobleaching phenotype (Figure 4a) and sucrose indirectly promotes *POR* gene expression (Mu *et al*., 2022), we quantified the transcript levels of *PORA* and *PORB* in etiolated seedlings grown with or without 2% sucrose supplementation. As demonstrated in Figure 6b, the transcript levels of *PORA* and *PORB* in the two *abi4* mutants were similar to those in the wild type, irrespective of the presence or absence of sucrose. In fact, the two *abi4* mutants resembled the *35Spro*::*GLK2*-*GFP* line in terms of having slightly higher transcript levels of both *POR*s compared to those in the wild type. Therefore, we conclude that the photobleaching observed in etiolated *abi4* mutant seedlings is not a result of low *POR* transcript levels.

To investigate the cause of Pchlide overaccumulation in etiolated *abi4* mutants, we quantified the transcript levels of three main GLK target genes that are involved in Pchlide biosynthesis. As seen in Figure 6c, both *HEMA1* and *CHLH* transcripts accumulated to slightly higher levels in the *35Spro*::*GLK2*-*GFP* line and the two *abi4* mutants compared to the wild type, although ANOVA and Tukey’s post hoc tests did not indicate statistically significant differences. *GUN4* transcript level was significantly upregulated in the *35Spro*::*GLK2*-*GFP* line, and its transcript levels in the two *abi4* mutants were slightly elevated from the wild-type level, albeit not significantly. In comparison, the transcript levels of all three genes were lowest in *glk1 glk2* and *glk1 glk2 abi4*. Together, the data demonstrate that GLKs can at least partially promote the accumulation of *HEMA1*, *CHLH* and *GUN4* transcripts in darkness, and the transcript accumulation patterns of these three genes may explain to a limited degree the Pchlide overaccumulation in etiolated *abi4* mutant seedlings.

An overexpression of *LHCB*—encoding the Light Harvesting Complex proteins surrounding photosystem II—has been reported to exacerbate cotyledon photobleaching (Liu *et al*., 2018). Since *LHC*s are also a main class of GLK targets (Waters *et al*., 2009), we quantified their transcript levels in 5-day-old etiolated seedlings. Amongst the *LHC*s we examined, five had a higher transcript level in the *35Spro*::*GLK2*-*GFP* line than in the wild type: *LHCB2.1*, *LHCB2.2*, *LHCB2.3*, *LHCB3* and *LHCB6* (Figure 7a). Their transcript levels were also either significantly or slightly elevated, in the two *abi4* mutants. Importantly, the accumulation levels of these transcripts were again lowest in *glk1 glk2* and *glk1 glk2 abi4* mutants, showing that the high *LHC* induction in etiolated *abi4*-1 mutant seedlings was completely dependent on *GLK*s. Furthermore, because *LHCB* overexpression impairs cotyledon greening in the light by promoting precocious prothylakoid (PT) proliferation in darkness without affecting Pchlide accumulation (Liu *et al*., 2018), we examined the etioplast ultrastructure in these 5-day-old etiolated seedlings using transmission electron microscopy (TEM). As shown in Figure 7b, we observed that the membrane structures within some etioplasts showed an “inverted staining”,

**Figure 7.**
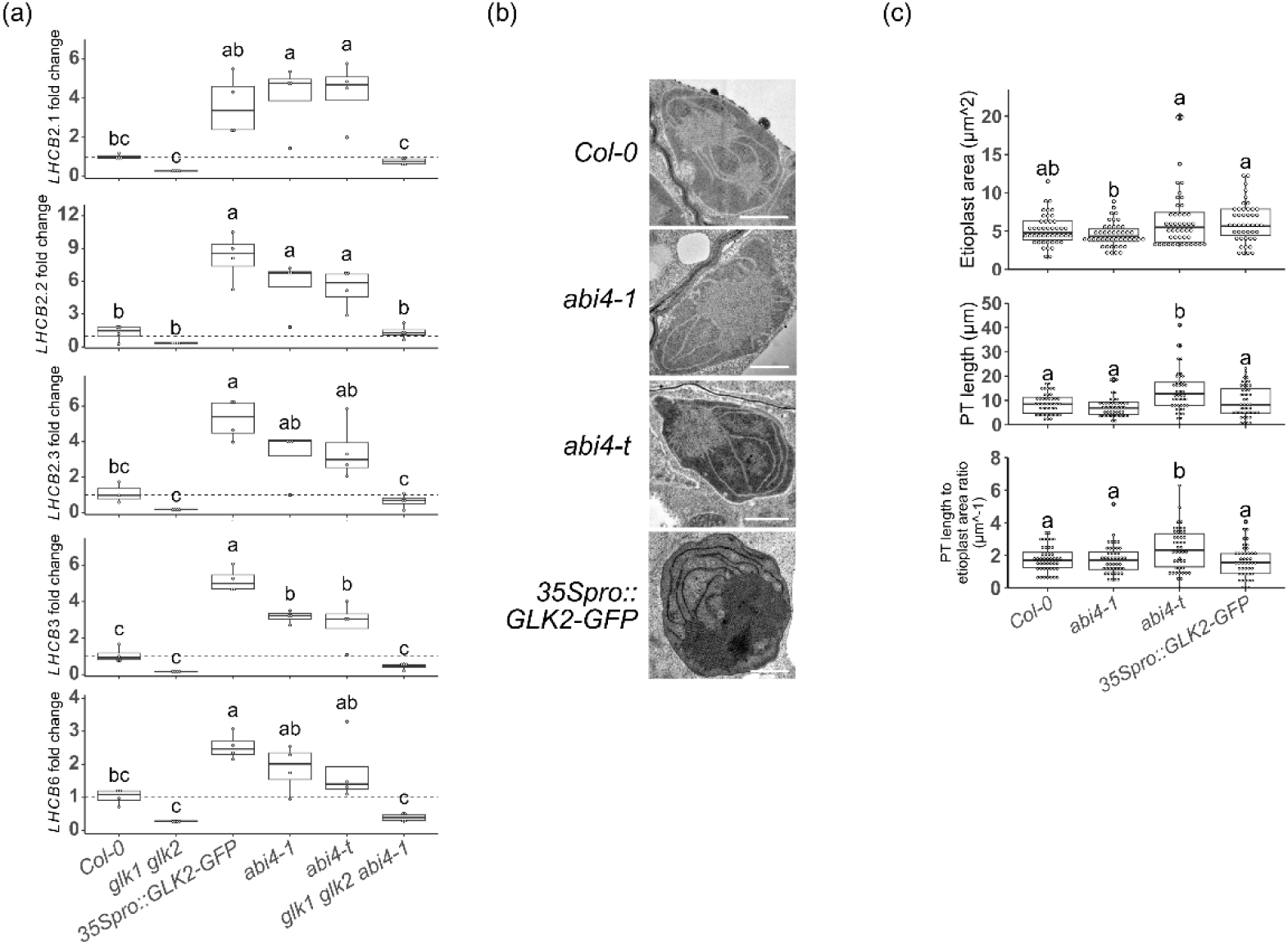
Seedlings of *abi4* mutants sometimes have over-differentiated etioplasts, **(a)** Five *LHCB*genes show high transcript levels in etiolated *abi4* mutants seedlings. Seedlings were grown on MS agar plates in darkness for five days without sucrose, and the transcript levels were quantified using RT-qPCR. Mean transcript level in *Col-0* was set to 1 (indicated by the dashed line), and data in each panel were compared with ANOVA and Tukey’s post hoc tests. Genotypes which are significantly different are labelled with different letters (n=4 replicate extracts), **(b)** Representative images of the etioplast ultrastructure in 5-day-old etiolated cotyledons under transmission electron microscope, scale bars = 1 µm. **(c)** Quantification of etioplast area (upper panel), prothylakoid (PT) length (middle panel) and the PT length to etioplast area ratio (lower panel). Genotypes which are significantly different are labelled with different letters (ANOVA and Tukey’s post hoc tests, n=45 etioplasts).

where both the PLB and the PTs appeared as white instead of black against the dark stroma. The occurrence of such inverted staining did not strictly correlate to certain genotypes, since seedlings of the *Col-0* and *abi4-t* genotypes contained both etioplasts that were stained normally and etioplasts with inverted staining. Nevertheless, the staining status *per se* did not affect the etioplast ultrastructure. When compared with the wild type, the *abi4* mutants and the *35Spro::GLK2-GFP* seedlings did not have significantly different etioplast sizes (Figure 7c). However, the *abi4-t* mutant etioplasts showed significantly longer PT than the wild type, and this result remained the same when the PT length was adjusted to the total etioplast area (Figure 7c). Overall, our data indicate that the accumulation patterns of *LHCB* transcripts in darkness may cause precocious etioplast-to-chloroplast differentiation, which would partially contribute to the greening-related phenotypes of etiolated *abi4* mutant seedlings, and these data hint at the possibility that these mutant seedlings have a higher activity of GLKs.

Lastly, since ABI4 physically interact with GLK1 and GLK2, we hypothesized that chromatin immunoprecipitation (CHIP) may reveal certain regulatory regions of GLK target genes as physically interacting with ABI4 proteins. By examining previously published CHIP datasets acquired using transgenic Arabidopsis grown in various conditions (Waters *et al*., 2009; Tu *et al*., 2022; Reeves *et al*., 2011; Yu *et al*., 2016; Liu *et al*., 2020), we discovered that both ABI4 and either GLK1 or GLK2 bind to the regulatory regions of *LHCB2.2*, *LHCB2.4*, *LHCB4.2*, *LHCB6* and *CHLH*. Together, these CHIP data corroborate the physical and functional ABI4-GLK interaction that we propose here.

## Discussion

In the present work, we identified ABI4 as a GLK-interacting protein in yeast two-hybrid (Figure 1) and *in planta* (Figure 2). ABI4 belongs to the AP2/ERF transcription factor family and is well-conserved in land plants (Gregorio *et al*., 2014). Arabidopsis loss-of-function *abi4* mutants were first identified based on their insensitivity to ABA during seed germination (Finkelstein, 1994; Finkelstein *et al*., 1998). ABI4 participates in additional signalling pathways, including responses to sugars and multiple hormones, while its role in retrograde signalling from the chloroplast to the nucleus has been controversial (Chandrasekaran *et al*., 2020).

We report here that etiolated, dark-grown seedlings of two *abi4* mutants are phenotypically similar to seedlings overexpressing GLK2 (*35Spro*::*GLK2*-*GFP*): etiolated *abi4* mutant and *35Spro*::*GLK2*-*GFP* seedlings contain more Pchlide (Figure 3), fail to green when exposed to light (Figure 4), generate more ^1^O_2_ after the dark-to-light transition (Figure 5), accumulate higher transcript levels of *LHCB*s in darkness (Figure 7) than the wild type. This suggests that GLKs and ABI4 have antagonistic functions. Indeed, *glk1 glk2* mutant seedlings have opposite phenotypes to those of *abi4* mutants: etiolated *glk1 glk2* mutant seedlings contain less Pchlide (Figure 3), green normally when exposed to light (Figure 4), generate low ^1^O_2_ upon light exposure (Figure 5) and accumulate slightly lower transcript levels of *PhANG*s in darkness (Figure 6 & 7) than the wild type. The direct protein-protein interaction between GLKs and ABI4 indicates that the antagonistic activities of GLKs and ABI4 are a result of the GLK-ABI4 interaction. Our finding that the *abi4* mutant phenotype fully depends on *GLK1* and *GLK2* supports this idea and suggests that ABI4 acts upstream of GLK1 and GLK2 by inhibiting the activities of the GLK1 and GLK2 proteins. The alternative possibility that ABI4 represses the expression of *GLK1* and *GLK2* can be excluded since *GLK1* and *GLK2* transcript levels were unaltered in *abi4* mutants.

### GLK and ABI4 antagonistically impact cotyledon greening due to their effects on Pchlide and singlet oxygen accumulation

In our greening assays, etiolated *abi4*-*1* and *abi4*-*t* mutant seedlings exhibited reduced greening rates compared to the wild type after the seedlings were exposed to light (Figure 4). This is not due to any major defect in chlorophyll biosynthesis, because *abi4* mutant seedlings green normally when grown without prior etiolation (Figures S1 & 4c) or even after 2 days of etiolation (Figure 4a). This is confirmed by Shu et al. (2013) where both *abi4*-*1* and *abi4*-*t* mutant seedlings achieved similar greening percentages compared to *Col*-*0* when grown with 1% sucrose and without prior etiolation. Since Pchlide is a photosensitizer that can inhibit seedling greening in light (Meskauskiene *et al*., 2001), we hypothesized that the photobleaching of etiolated *abi4* mutant seedlings is due to an overaccumulation of Pchlide in these seedlings. We found that etiolated *abi4* mutant seedlings, as well as etiolated *35Spro*::*GLK2*-*GFP* seedlings, accumulated more Pchlide than wild-type seedlings (Figure 3), which correlated with lower greening rates after seedlings were exposed to light (Figure 4). In contrast, the *glk1 glk2* mutant accumulated less total Pchlide than the wild type as etiolated seedlings (Figures 3a & 3b), and it showed high greening rates in the cotyledon greening assay (Figure 4). Importantly, the *glk1 glk2* mutations returned both the Pchlide accumulation (Figure 3) and the cotyledon greening rate (Figure 4) of *abi4-1* mutant seedlings to a wild-type level. We therefore conclude that the Pchlide overaccumulation in etiolated *abi4* mutant seedlings causes the photobleaching of these seedlings after the dark-to-light transition, and this Pchlide overaccumulation may be due to de-repressed GLK activity.

Next, we hypothesized that the photobleaching of etiolated *abi4* mutant seedlings in light is caused by singlet oxygen generated by Pchlide photosensitization. Singlet oxygen in plants is primarily generated in the plastids, either by chlorophyll molecules during photosynthesis or by chlorophyll precursors during dark-to-light transitions (Wang *et al*., 2020). ^1^O_2_ is toxic to the cell (Apel and Hirt, 2004), and its accumulation with other ROS species can lead to programmed cell death (Wagner *et al*., 2004; Kim *et al*., 2012). Because of the direct role of GLKs in promoting Pchlide biosynthesis, it was not surprising to find that etiolated *glk1 glk2* seedlings contained less ^1^O_2_ and showed a high greening rate, while etiolated *35Spro*::*GLK2-GFP* seedlings contained more ^1^O_2_ and showed a low greening rate upon light exposure compared to the wild type (Figures 4 & 5). A recent publication also supports the conclusion that limiting GLK activity in dark-grown seedlings acts to promote cotyledon greening during de-etiolation (Tachibana *et al*., 2025). We also observed high ^1^O_2_ levels and low greening rates in both *abi4* mutants after their etiolated seedlings were transferred to light (Figures 4 & 5). This seems to contrast with a previous study in which two *abi4* loss-of-function mutations both rescued the lethal phenotype of the *flu* mutant under dark/light cycles (Zhang *et al*., 2023). The *flu* mutant seedlings accumulate at least four times as much Pchlide as the wild type during etiolation, leading to rapid cotyledon bleaching and death after the dark-to-light transition (Zhang *et al*., 2023; Meskauskiene *et al*., 2001). Intriguingly, the *abi4* mutations did not alter the Pchlide accumulation in etiolated *flu* mutant seedlings, nor the ^1^O_2_ accumulation upon light exposure, but they lowered the transcript levels of ^1^O_2_-responsive genes in light (Zhang *et al*., 2023). Since chloroplast ^1^O_2_ stress responses are controlled by multiple pathways (Tano *et al*., 2023), we suggest the pathways mediating the photobleaching of etiolated *flu* and *abi4* mutant seedlings in light may be different.

We further hypothesized that etiolated *abi4* mutant seedlings accumulate high levels of Pchlide due to an upregulated Pchlide biosynthetic pathway. We showed that the *HEMA1*, *CHLH* and *GUN4* transcript levels in the etiolated seedlings of the *35Spro*::*GLK2*-*GFP* line and the two *abi4* mutants were at least slightly elevated compared to the wild type (Figure 6c). These transcripts possibly upregulated the Pchlide biosynthesis in the seedlings. The similarity between etiolated *35Spro*::*GLK2*-*GFP* and *abi4* mutant seedlings suggests again that GLK protein activity may be de-repressed in etiolated *abi4* mutant seedlings. *HEMA1*, *CHLH* and *GUN4* are activated by GLKs in light-grown plants (Waters *et al*., 2009), and we demonstrated here that their transcript accumulation in etiolated seedlings in the *abi4-1* mutant background is dependent on *GLK1* and *GLK2* (Figure 6c). As expected, these *GLK*-dependent transcript levels correlated with the *GLK*-dependent Pchlide accumulation during etiolation (Figures 3a & 3b) and greening patterns after the dark-to-light transition (Figure 4). Since the transcript levels presented here came from end-point analyses, future studies can generate transgenic Arabidopsis lines each expressing a luciferase driven by promoters of *HEMA1*, *CHLH* or *GUN4*, whereby the transcriptional dynamics of these promoters can be monitored in real-time by luciferase luminescence during etiolation and compared among different genotypes. Additional GLK and ABI4 target genes should be identified in order to fully explain the Pchlide accumulation pattern observed here.

We also investigated whether the *POR* transcript level is altered in etiolated *abi4* mutant seedlings, due to the direct role of PORs in Pchlide photoreduction. Knocking out either *PORA* or *PORB* produces seedlings with smaller PLBs, unstructured PTs and a delayed-greening phenotype (Masuda *et al*., 2009; Masuda *et al*., 2003). *PORA* and *PORB* are both transcriptionally activated by GLKs in seedlings grown under long-day photoperiod (Waters *et al*., 2009). In comparison, we observed only a slight transcript overaccumulation of either *POR*s in etiolated *35Spro*::*GLK2*-*GFP* seedlings compared to the wild type (Figure 6b). Moreover, we observed no significant change in *PORA* and *PORB* transcript levels in both *abi4* mutants compared to the wild-type etiolated seedlings (Figure 6b). This differs from the report of Xu et al. (2016), where etiolated *abi4-1* mutant seedlings were shown to contain a lower *PORA* transcript level compared to that in the wild type. Our observation remained the same, either without supplementary sucrose in the growth medium, or with 2% supplementary sucrose, as that used by Xu et al. (2016). We therefore conclude that the *POR* transcript level is unlikely to have significantly affected our results.

### Impact of GLKs and ABI4 on the expression of *PhANG*s and on the etioplast structure

Beyond the PLB, the ultrastructure of the PTs also affects the greening process (Li *et al*., 2024; Liu *et al*., 2018). Overexpressing either *LHCB1.1* or *LHCB2.1* produces seedlings with low greening rates when the seedlings are exposed to light after prolonged etiolation (Liu *et al*., 2018). These transgenic seedlings contain a wild-type level of total Pchlide, but they develop etioplasts with no PLBs and only over-proliferated PTs. This agrees with our findings that the *35Spro*::*GLK2*-*GFP* line and the two *abi4* mutants, which accumulated more *LHCB* transcripts than the wild type as etiolated seedlings (Figure 7a), showed lower greening rates in the cotyledon greening assay (Figure 4). In seedlings of these genotypes, only the *abi4-t* mutant exhibited over-proliferated PTs compared to the wild type (Figure 7b & 7c). This could be due to the fact that TEM sections only represent the three-dimensional etioplast ultrastructure at a two-dimensional level, and that etioplasts show high structural variation. Amongst the different LHC isoforms, at least LHCB2 accumulates during seedling etiolation to a level detectable by western blotting (Armarego-Marriott *et al*., 2019), therefore, future studies can investigate whether the *35Spro*::*GLK2*-*GFP* line and the two *abi4* mutants accumulate more LHCB proteins than the wild type in etiolated seedlings. Additionally, we observed that some etioplasts showed an “inverted staining” (Figure 7b). Although such inverted staining has occurred in other etioplast studies before (Liang *et al*., 2022; Vinti *et al*., 2005), we presently cannot explain why in our experiment even seedlings of the same genotype were found to contain both normally stained and invertedly stained etioplasts.

### Model on the antagonistic activities of GLKs and ABI4

The present study suggests that the ABI4-GLK protein-protein interaction regulates the greening of etiolated seedlings in light, and this agrees with previous reports where other protein-protein interactions between nuclear proteins regulate Pchlide biosynthesis and etioplast development (Chen *et al*., 2013; Liu *et al*., 2018; Sun *et al*., 2019). At the moment, the mechanism(s) by which ABI4 represses GLK activity is not clear, and in addition to darkness, this repression could occur in other conditions where limiting *PhANG*s expression becomes necessary. ABI4 may reduce GLK protein stability, inhibit GLK dimerization, limit GLK binding to its target promoters and/or repress the transcriptional activation capacity of GLK. D. Zhang et al. (2021) reported that GLK1 and GLK2 are protected from proteasomal degradation after being phosphorylated by BRASSINOSTEROID INSENSITIVE 2 (BIN2), and since BIN2 is inactivated by darkness, GLK proteins are less stable in darkness. N. Kim et al. (2023) revealed that REPRESSOR OF PHOTOSYNTHETIC GENES 1 (RPGE1) represses GLK1 homodimerization and DNA-binding, thereby limiting *PhANG*s activation in shade. Intriguingly, it was shown by M. Li et al. (2022) that treating plants with salicylic acid alters the GLK interactome, reduces the binding of GLK to *PhANG*s promoters, and enhances the stress response. These reports illustrate the various mechanisms whereby GLK protein activity can be modulated.

Since ABI4 physically interacts with GLK, a direct binding of ABI4 to GLK target promoters is not necessary for ABI4 to inhibit GLK target gene expression. Nevertheless, ABI4 has been shown to bind *LHCB* promoters in Y1H, *in vitro* and *in planta* (Koussevitzky *et al*., 2007; Guo *et al*., 2016; Zhang *et al*., 2023). We therefore cannot exclude the possibility that ABI4 can regulate the expression of certain *PhANG*s independently of GLKs, or that ABI4 regulates GLK functions in light, similar to how PSEUDO-ETIOLATION IN LIGHT (PEL) proteins suppress GLK activities (Han *et al*., 2024). Nonetheless, the antagonistic GLK-ABI4 interaction that we propose here may integrate another layer of complexity to a transcriptional factor network based on direct protein-protein interactions, including those between GLKs and HY5 (Zhang *et al*., 2024) and those between ABI4 and PIF4 (Luo *et al*., 2024; Song *et al*., 2024). The co-action of these transcriptional factors would serve to fine-tune the pigment biosynthesis and plastid development according to the needs of the developing seedling.

## Materials and methods

### Plant material and growth conditions

All Arabidopsis lines used in this study were in the Columbia ecotype, and we list their NASC stock IDs here. Both the *glk1 glk2* mutant (N9807) and the *35Spro*::*GLK2*-*GFP* line (N2107721) were described previously (Fitter *et al*., 2002; Waters *et al*., 2008). The mutants *abi4-1* (N73626), *gl1* (N1688), *abi4-103 gl1* (N3838) and *abi4-t* (N580095) were described in the text.

Seeds were surface-sterilized, stratified for 3 days in darkness at 4°C, then sown on Murashige and Skoog (MS) media containing 10 g/L Agar (SERVA) (without sucrose unless indicated otherwise). The sown seeds were exposed to white light (100 μmol m-^2^ s^-1^) for 4 hours to induce germination, then left in a growth chamber either under the same white light or in darkness at 21°C. For analysis of chlorophyll content, plants were grown on soil in a climate-controlled walk-in growth chamber under long-day photoperiod (16 h of 100 μmol m-^2^ s^-1^ white light/8 h darkness) at 21 °C.

### Generation and verification of the *glk1 glk2 abi4-1* mutant

The *glk1 glk2 abi4-1* triple mutant was generated by crossing *glk1 glk2* with *abi4-1*. From the F2 population, those homozygous for *glk1* and *glk2* were identified by their pale-green cotyledons. Genomic DNA was isolated individually from these plants and subjected to a dCAPS analysis for *abi4-1* homozygosity. A sequence encompassing the *abi4-1* locus was PCR-amplified, and the PCR products were digested by AluI. The sequences of primers used for dCAPS analysis are listed in Supplementary Table S1.

### Cloning of constructs

For Y2H, we generated the entry clones for BD-GLK1_Δ1-154,_ BD-GLK2_Δ1-148_, BD-ABI4_Δ188-289_, AD-GLK1, AD-GLK2 by first PCR-amplifying the coding sequences from Arabidopsis cDNA and cloning into pENTR3C (Thermo Fisher) via homologous recombination (ClonExpress, Vazyme). The entry clone for AD-ABI4 was obtained by isolating the pDEST22-ABI4 vector from the REGIA Y2H library (Paz-Ares *et al*., 2002) and mixing pDEST22-ABI4 with pDONR221 (Thermo Fisher) in a BP Gateway reaction (Thermo Fisher). Then the destination vectors were each cloned by mixing the respective entry clone with a GAL4-based BD- or AD-vector (pAS2-GW and pACT2-GW) (Clontech) in an LR Gateway reaction (Thermo Fisher).

For FRET-FLIM assays, pAMARENA (Steffens *et al*., 2014) and pENSG-YFP (Laubinger *et al*., 2006) were used for N-terminal tagging with mCherry and YFP, respectively. The destination vectors were cloned via LR reactions using either the entry clone pENTR3C-GLK or pDONR221-ABI4.

For the luciferase complementation imaging assay, all nLUC- and cLUC-expression vectors were generated by conventional digestion-ligation cloning. GLK1 and GLK2 were cloned into the pCambia1300-nLUC vector, and ABI4 was cloned into pCambia1300-cLUC (Chen *et al*., 2008), in all cases using the KpnI and SalI restriction sites. The sequences of primers used for cloning are listed in Supplementary Table S1.

### Yeast two-hybrid assays

Gal4-based yeast two-hybrid assays, specifically the histidine growth selection assay, were performed. The BD- and AD-plasmids were transformed into the yeast strain AH109 (Clontech) using the Frozen-EZ Yeast Transformation II Kit (Zymo Research). Transformants were then selected on Synthetic Drop-out auxotrophic plates (SD -L -W) (Takara Bio) containing 2% glucose. 10 co-transformed yeast colonies were suspended in sterile water and plated on both SD -L -W and SD -L -W -H plates containing 2% glucose. The yeast was grown at 30 °C, and 3-aminotriazole (3-AT) was used in the SD -L -W -H plates to prevent autoactivation.

### FRET-FLIM assays

For the FRET-FLIM assays, leek epidermal cells were transiently transformed with different FRET constructs by particle bombardment as described by Ponnu et al. (2019). The FLIM experiments were also performed according to (Ponnu *et al*., 2019), with the difference that the curve-fitting processes were done in the software SymPhoTime 64 (PicoQuant).

### Firefly luciferase complementation (LCI) imaging assay

Constructs expressing GLK1, GLK2 or ABI4 proteins fused to the C-terminal or N-terminal half of firefly luciferase were transformed into the *Agrobacterium tumefaciens* strain *GV3101::pMP90*, and different combinations of transformed *Agrobacterium* were infiltrated into tobacco leaves. The infiltration procedure and luminescence imaging were performed as described in (Kreiss *et al*., 2023). The luminescence intensity was quantified, and the bright-field and luminescence images were overlapped using Fiji (Schindelin *et al*., 2012). Data were plotted without normalization.

### Pchlide, Chlide and chlorophyll quantification

For Pchlide and Chlide quantification, 15 etiolated seedlings per replicate were harvested from MS agar plates in darkness. The samples were lysed and homogenized in 300 μl ice-cold 80% acetone, then centrifuged at 5,000 g for 2 min. 100 μl of the supernatant was aliquoted into a white 96-well plate. The fluorescence emission spectra of the sample were scanned from 600 to 700 nm using a spectrophotometer with an excitation wavelength of 440 nm (Tecan Infinite 200 PRO).

For chlorophyll quantification of plants grown in long-day photoperiod, 50-100 mg of leaf tissue per replicate was harvested and weighed, frozen in liquid nitrogen and ground to fine powder with a TissueLyser (Qiagen). 1.5 ml 80% acetone was added and samples were vortexed for 5 min then centrifuged at 13.200 rpm for 5 min. Finally, 100 μl of the supernatant was aliquoted into a transparent 96-well plate, and the absorbance was measured at 645 and 663 nm (Tecan Infinite 200 PRO). Total chlorophyll in mg per g fresh weight is calculated using the following formula: ((A663*0.00802)+(A645*0.0202))×1.5/fresh weight (Arnon, 1949). For chlorophyll quantification in the cotyledon greening assay, the procedure was kept the same, with the difference that 25 representative seedlings per replicate were harvested, frozen, ground, then added to 150 µl 80% acetone. Total chlorophyll in ng per seedling is calculated using the following formula: ((A663*8.02)+(A645*20.2))×0.15/25.

### Cotyledon greening assays

Around 100 seeds sown on each replicate MS agar plate were stratified, grown for varied numbers of days in the dark, followed by 3 d in Wc (100 μmol m^-2^ s^-1^), all at 21°C. After 3 days, the germinated seedlings were classified into green/bleached by manually examining the phenotype of individual seedlings.

### Fluorescence imaging of SOSG staining

Seedlings were incubated with 100 μM SOSG (Lumiprobe) in the dark for 20 min. The excess SOSG solution was then removed, and the seedlings were washed with sterile water. The SOSG signal was captured by laser excitation at 488 nm and emission at 525 nm. Chlorophyll autofluorescence was visualized by excitation at 610 nm and emission at 630 nm. The laser and detector settings were kept the same for all measurements. The imaging was performed using the Leica TCS SP8 (Leica microsystems). Image analyses were performed in Fiji (Schindelin *et al*., 2012).

### Reverse transcription-quantitative PCR (RT-qPCR)

Total RNA was extracted from Arabidopsis seedlings (around 40 seedlings per replicate), using the NucleoSpin RNA Plant Mini Kit (Macherey Nagel) according to the manufacturer’s instructions. For cDNA synthesis, 1 µg RNA, 2 µl 10 µM Oligo-dT primer and RNase-free water were added to a total volume of 27 µl. The reaction was incubated at 65°C for 5 minutes, then cooled on ice for 2 minutes. Subsequently, 8 µl 5 x RevertAid buffer, 4 µl 10 mM dNTPs and 1 µl RevertAid H Minus Reverse Transcriptase (Thermo Fisher) were added to the reaction, and the mix was incubated at 37°C for 5 minutes, at 42°C for 1 h for cDNA synthesis, then at 70°C for 5 minutes for heat-inactivation.

For RT-qPCR analysis, 0.5 µl of cDNA was mixed with 0.5 µl 10 µM forward and reverse primers and 5 µl GoTaq qPCR Master Mix (Promega) in a 10 µl reaction. The RT-qPCR was performed on a QuantStudio 5 Real-Time PCR System (Thermo Fisher). The relative transcript levels were calculated using the 2^−ΔΔCt^ method (Livak and Schmittgen, 2001). The sequences of primers used for qPCR are listed in Supplementary Table S1.

### Transmission electron microscopy

Seeds sown on MS agar plates were first stratified, then kept in darkness at 21 °C for 5 days. On Day 1, whole seedlings (∼30 per genotype) were fixed in 2.5% glutaraldehyde + 4% PFA in 0.1 M phosphate buffer (pH 7.4) by vacuum infiltration for 1 h under a fume hood and stored overnight at 4 °C. On Day 2, samples were washed twice quickly and then three times for 5 min each with 0.1M phosphate buffer (pH 7.4) at room temperature. Post-fixation was carried out in osmium solution (1% OsO_4_, 2 mM CaCl_2_, 0.8% potassium ferricyanide in 0.1 M phosphate buffer pH 7.4) for 1 h in the dark on ice. Next, samples were washed seven times with 0.1 M phosphate buffer (pH 7.4) at RT. Seedlings were then dehydrated in 50% ethanol (1 h on ice), and subsequently in 70% ethanol (overnight at 4°C). On Day 3, *en bloc* staining was performed by incubating samples in 1% uranyl acetate in 70% ethanol for 1h in the dark at RT, then seedlings were washed twice quickly and then five times 5 min each in 70% ethanol on ice. Dehydration was carried out by graded ethanol series (80%, 90%, 96%, and 100%) for 30 min at each step (100% twice) on ice. Afterwards, the samples were washed with 100% acetone (2x30 min), followed by overnight rotation in in an Epon^TM^:acetone (1:3) mixture (Merck, Epoxy Embedding Medium kit, cat. no. 45359-1EA-F) at RT. On Day 4, Epon^TM^ infiltration was continued by rotating samples in progressively higher concentrations of Epon™:acetone (1:2 for 30 min, 1:1 for 1 h, 2:1 for 1 h, and 3:1 for 2 × 1 h), followed by incubation in pure Epon™ for 2 h. Fresh Epon™ was then added for the overnight rotation at RT. On Day 5, seedlings were incubated in fresh Epon^TM^ for 2 h at RT, then the cotyledons were excised and embedded in moulds filled with Epon^TM^, which were polymerized at 65°C for 72h. Ultrathin sections (70 nm) were prepared using an ultramicrotome (UC Enuity, Leica Microsystems). Sections were stained with 2% aqueous uranyl acetate for 5 min, rinsed with distilled water, and subsequently stained with Reynolds’ lead citrate (Reynolds, 1963) for 4 min under CO₂-free conditions, followed by a final rinse with distilled water. The sections were examined with a transmission electron microscope (Zeiss EM 902) operated at 80 keV, and images were captured using a slow-scan CCD camera (Type 7899). Three seedlings per genotypes were examined.

### Statistical analyses

All statistical analyses were performed in the R environment (R Core Team, 2019), and the data were visualized using the ggplot2 package (Wickham, 2009). Each individual replicate is plotted as a dot, and summary statistics are plotted as indicated in each figure. For boxplots, the hinges on each box indicates the lower and upper quantiles, and the whiskers show data range. The median is represented by a horizontal line within the box. No data were excluded from the analyses.

### Accession numbers

The Arabidopsis genes included in this study can be identified in The Arabidopsis Information Resource (www.arabidopsis.org) under these accession numbers: *ABI4* (AT2G40220), *GLK1* (AT2G20570), *GLK2* (AT5G44190), *PORA* (AT5G54190), *PORB* (AT4G27440), *HEMA1* (AT1G58290), *CHLH* (AT5G13630), *GUN4* (AT3G59400), *LHCB2*.1 (AT2G05100), *LHCB2*.*2* (AT2G05070), *LHCB2*.*3* (AT3G27690), *LHCB3* (AT5G54270), *LHCB6* (AT1G15820).

## Author contributions

PY and UH conceived the project and wrote the article. All authors read and revised the article. PY and FS performed the physiological and molecular experiments. PY and MB performed the TEM experiment. All authors analyzed the data.

## Acknowledgements

We thank Dr. Kim Boutilier for providing the *abi4-1* seeds, and the Nottingham Arabidopsis Stock Centre (NASC) for providing the other mutant and transgenic lines. We are also grateful to the greenhouse staff at the Biocenter for cultivating our plants.

## Supporting Information

Figure S1. Chlorophyll accumulation is not altered in light-grown *abi4* mutants.

Figure S2. Verification of the *glk1 glk2 abi4-1* triple mutant.

Table S1. Primers used in this study.

## Data availability statement

The data that support the findings of this study are available in the supplementary material of this article.

