## Supplemental figures for "Arabidopsis GLK transcription factors interact with ABI4 to modulate cotyledon greening in light-exposed etiolated seedlings"

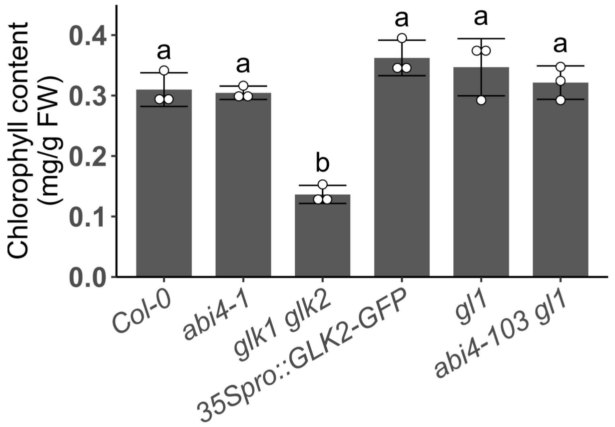


Figure S1. Chlorophyll accumulation is not altered in light-grown *abi4* mutants. Seedlings are grown for 10 days on soil in a long-day phytochamber with 100 µmol photons $m^{-2}$ $s^{-1}$ white light. Data were normalized to per gram fresh weight (/g FW). Genotypes which are significantly different are labelled with different letters (n=3 replicate extracts, ANOVA and Tukey's post hoc tests), error bars indicate SD.

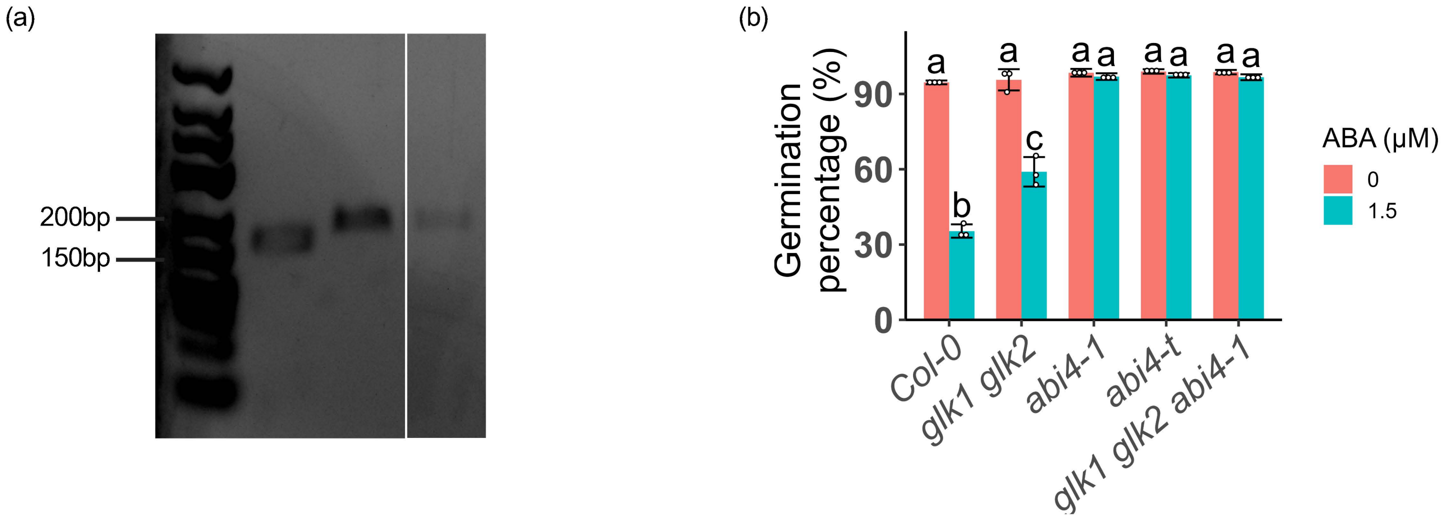


Figure S2. Verification of the *glk1 glk2 abi4-1* triple mutant. (a) Genotyping by dCAPS, samples were ran on a 3% agarose gel at 100 V for 25 min. From left to right: GeneRuler low range DNA ladder; sample with *Col-0* gDNA; sample with *abi4-1* gDNA; sample with homozygous *glk1 glk2 abi4-1* gDNA. The *Col-0* fragment is predicted to be 25 bp smaller than the *abi4-1* fragment. (b) Germination phenotype on MS agar plates ± ABA. Genotypes which are significantly different are labelled with different letters (n=3 replicate plates, ANOVA and Tukey's post hoc tests), error bars indicate SD.
