## Supplemental Table 1 for "Arabidopsis GLK transcription factors interact with ABI4 to modulate cotyledon greening in light-exposed etiolated seedlings"

Supplementary Table S1. Primers used in this study

| | |
|---|---|
| Primers used for dCAPS | |
| abi4-1-genotyping-AluI-F | CTCAACTTAACCCCTTCGTCTCCTTC |
| abi4-1-genotyping-AluI-R | GGGATACCGTACGGACCAAAGTTAG |
| Primers used for cloning Y2H constructs | |
| pENTR3C-GLK1-F | ggatccggtaccgaattcATGTTAGCTCTGTCTCCGGC |
| pENTR3C-GLK1-R | gagtgcggccgcgaattcTCAGGCACAAGACGCGG |
| pENTR3C-GLK2-F | ggatccggtaccgaattcATGTTAACTGTTTCTCCGGCTC |
| pENTR3C-GLK2-R | gagtgcggccgcgaattcTCAAGGAAGAGGAGGAACAT |
| pENTR3C-GLK1$_{\Delta 1-154}$-F | ACTGGATCCGGTACCGAATTcAAAGTGGATTGGACACCAGAGC |
| pENTR3C-GLK1$_{\Delta 1-154}$-R | CTCGAGTGCGGCCGCGAATTTCAGGCACAAGACGCGGTCG |
| pENTR3C-GLK2$_{\Delta 1-148}$-F | ACTGGATCCGGTACCGAATTCAAGGTGGATTGGACGCCGGA |
| pENTR3C-GLK2$_{\Delta 1-148}$-R | CTCGAGTGCGGCCGCGAATTTCAAGGAAGAGGAGGAACATTAGAAA |
| pENTR3C-ABI4$_{\Delta 188-289}$-F | ACTGGATCCGGTACCGAATTcATGGACCCTTTAGCTTCCCA |
| pENTR3C-ABI4$_{\Delta 188-289}$-R | CTCGAGTGCGGCCGCGAATTATAGAATTCCCCCAAGATGGGA |
| Primers used for cloning LCI constructs | |
| pCAMBIA1300-GLK1-nLUC-F | gacgagctcggtaccATGTTAGCTCTGTCTCCGGCG |
| pCAMBIA1300-GLK1-nLUC-R | gtacgagatctggtcgacGGCACAAGACGCGGTCG |

| | |
|---|---|
| pCAMBIA1300-GLK2-nLUC-F | ggggacgagctcggtaccATGTTAACTGTTTCTCCGGCTCCA |
| pCAMBIA1300-GLK2-nLUC-R | gcgtacgagatctggtcgacAGGAAGAGGAGGAACATTAGAAACTCC |
| pCambia1300-cLUC-ABI4-F | tacgcgtcccggggcggtaccATGGACCCTTTAGCTTCCCA |
| pCambia1300-cLUC-ABI4-R | acgaaagctctgcaggtcgacTTAATAGAATTCCCCCAAGATGG |
| Primers used for qPCR | |
| GLK1-F | CCGTATTTACCGACCGTAGCTACGAGA |
| GLK1-R | TACATCGTGTGATGCGGCGGCAGAG |
| GLK2-F | TGTGTGTAAGCAAGAGGGTGG |
| GLK2-R | CTACCCCTAATTGCTCCACCG |
| PORA-F | CCCTCTTCCCTCCTTTCCAG |
| PORA-R | GCTCCAATACACTCCCGACTTC |
| PORB-F | ACTTGCTCAGGTGGTGAGTG |
| PORB-R | CCACACTTTACGAGCCTTCT |
| HEMA1-F | GCTTCTTCTGATTCTGCGTC |
| HEMA1-R | GCTGTGTGAATACTAAGTCCAATC |
| CHLH-F | CATTGCTGACACTACAACTGC |
| CHLH-R | CTTCTCTATCTCACGAACTCCTTC |
| GUN4-F | CAATCTCACTTCGGACCAAC |
| GUN4-R | TTGAAACGGCAGATACGG |
| LHCB2.1-F | ACCCGGAGACATTCGCTAAGAAC |
| LHCB2.1-R | TGGATCAAGTTAGGGTTTCCGAGG |
| LHCB2.2-F | AGCAGATGGGCTATGTTGGGT |
| LHCB2.2-R | CTCAGCCAAGTTCAATGGGTCG |
| LHCB2.3-F | CGTCAAGTCTACTCCTCAGAGCATC |
| LHCB2.3-R | TAATATGCTTTGCGCGTGGATCAAG |
| LHCB3-F | GCCGGTTCACAAATCTTCTCCG |
| LHCB3-R | AGTAACTGGATCATCAGCGAGACC |
| LHCB6-F | TGGCTCGCTACCAGGAGATTTC |
| LHCB6-R | CAATTGGGTTCCAAGAAGCGACC |
